# Super-resolved polarisation-enhanced second harmonic generation for direct imaging of nanoscale changes in collagen architecture

**DOI:** 10.1101/2020.02.07.934000

**Authors:** Peter Johnson, Artemis Karvounis, H. Johnson Singh, Christopher J. Brereton, Konstantinos Bourdakos, Kerry Lunn, James JW Roberts, Donna E Davies, Otto L. Muskens, Mark G Jones, Sumeet Mahajan

## Abstract

Super-resolution (SR) optical microscopy has allowed the investigation of many biological structures below the diffraction limit, however, most of the techniques are hampered by the need for fluorescent labels. Non-linear label-free techniques such as Second Harmonic Generation (SHG) provide structurally specific contrast without the addition of exogenous labels, allowing observation of unperturbed biological systems. Here we achieve super-resolution SHG (SR-SHG) for the first time. We use the photonic nanojet (PNJ) phenomena to achieve a resolution of ~λ/6 with respect to the fundamental wavelength, a ~2.7-fold improvement over diffraction-limited SHG under the same imaging conditions. Crucially we find that the polarisation properties of excitation are maintained in a PNJ allowing the resolution to be further enhanced by detection of polarisation-resolved SHG (p-SHG) by observing anisotropy in signals. These new findings allowed us to visualise biological SHG-active structures such as collagen at an unprecedented and previously unresolvable spatial scale. Moreover, we demonstrate that the use of an array of self-assembled high-index spheres overcomes the issue of a limited field of view for such a method, allowing PNJ-assisted SR-SHG to be used over a large area. Dysregulation of collagen at the nanoscale occurs in many diseases and is an underlying cause in diseases such as lung fibrosis. Here we are able to demonstrate that pSR-SHG allows unprecedented observation of changes at the nanoscale that are invisible by conventional diffraction-limited SHG imaging. The ability to non-destructively image SHG-active biological structures without labels at the nanoscale with a relatively simple optical method heralds the promise of a new tool to understand biological phenomena and drive drug discovery.

## Introduction

The Abbe “diffraction limit” states that the resolution of an optical microscope is limited by its wavelength to ~λ/(2NA) with NA the numerical aperture of the medium. In biological microscopy, this poses a major limitation as a wealth of processes occur on a scale below the diffraction limit. Many techniques now exist to surpass this limit including STimulated Emission Depletion (STED) microscopy, localisation microscopies such as PALM/STORM and Structured Illumination Microscopy (SIM) collectively referred to as super-resolution (SR) techniques [1]. However, these techniques are either complex to implement or rely on the properties of fluorescent labels, or both. As such, achieving SR using label-free techniques such as SHG microscopy would be of great interest, but so far this modality has not seen the same success. SHG microscopy is a structurally specific technique that highlights non-centrosymmetric features without the addition of exogenous and potentially perturbative labels. SHG active materials include certain crystals such as Barium Titanate (BaTiO_3_) and biological macromolecules including the actin-myosin complex, microtubules and fibrillar collagen. The ability to perform label-free SHG imaging with sub diffraction-limited resolution therefore holds significant potential for biomedical research as changes in collagen structure and function underlie many diseases including fibroses and cancers.

To realise label-free SR-SHG imaging we have employed the photonic nanojet (PNJ) phenomena, as it does not rely on fluorescent switching/blinking and is simple to implement. A PNJ is a localised focus of light with an intensity cross section narrower than the diffraction limit that is able to propagate over multiple wavelengths [2]. Initial demonstrations of the phenomena used high refractive index spheres or cylinders to generate the PNJ and this is still the most common experimental implementation however, theoretical studies have investigated cuboids [3], cones, prism structures [4] and spheres with multiple/graded refractive indices [5]. Typically, dielectric spheres of sizes between 5 μm-100 μm in diameter are placed on the surface of a sample to focus the light into a PNJ, decreasing the illumination point spread function of the system [6]. Whispering gallery modes [7] and elements of solid immersion lens theory [8] [9] have also been suggested as contributing to the SR imaging capability of PNJ generating microspheres.

The focal spot of a PNJ, when generated using a microsphere is dependent on multiple parameters including sphere size, imaging wavelength and the refractive index contrast with the surrounding medium [10]. For biological imaging, the contrast has to be ideally maximised against the microscope immersion medium. Imaging with PNJs (schematic shown in Figure 1) has been typically performed in a widefield configuration, using both brightfield and epifluorescence [11,12]. It has also been demonstrated in a laser-scanning configuration [13] and therefore, is amenable to SHG microscopy.

**Figure 1.**
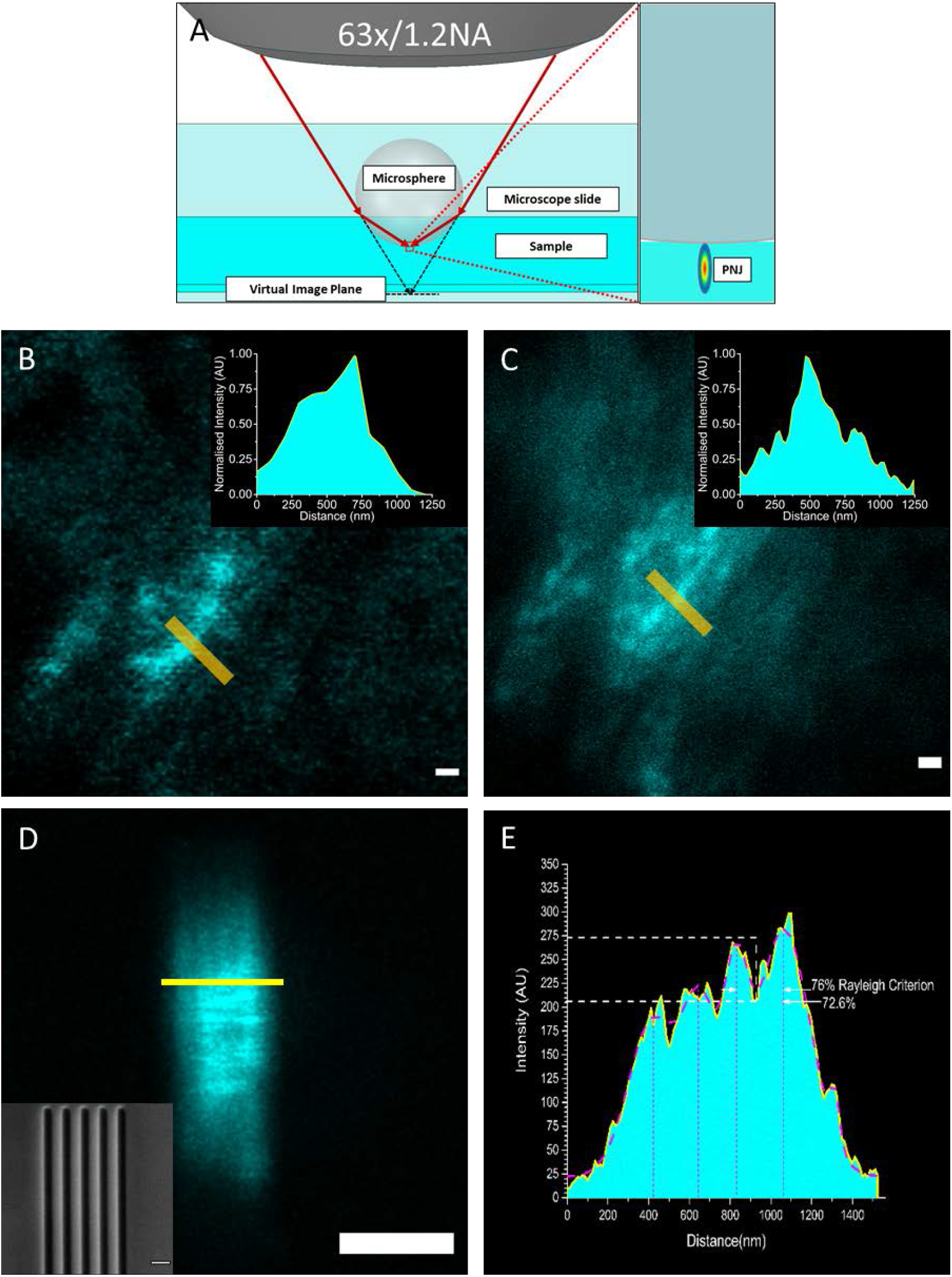
Establishing the limit of resolution of SR-SHG. **A:** A diagram depicting imaging using PNJs generated using a microsphere in a microscope setup. The microsphere is located on the surface of the sample and acts to focus the light to a sub-diffraction-limited spot. A super resolution virtual image is formed on the shadow side of the sphere. **B:** Diffraction limited SHG signal from a fibrotic lung tissue sample containing collagen. Scale bar 500 nm. **C:** SHG image of the same region taken using a PNJ. It is possible to resolve more fibres and their directionality. Insets show intensity plots taken over the yellow lines in B/C, More peaks are resolved in the PNJ image compared to the diffraction limited image. **D Inset** An SEM image of a the silicon sample, each dark slit is 100 nm wide and the slits are separated by 125 nm **D:** Image taken through a 14 µm sphere the silicon slit sample, the sub-diffraction limited features are resolved.. Scale bar 250 nm. **E**: An intensity plot taken across the yellow line in **D**. The plot shows 4 peaks corresponding to slits the peaks can be resolved with a drop to 72.6% intensity, satisfying the Rayleigh Criterion.

SHG interrogates the 2^nd^ order polarizability of materials and detects non-centrosymmetric structures. It is widely used to image fibrillar collagen in bioimaging [14]. SHG is polarisation sensitive; intensities depend on the relative orientation between the excitation and the SH active structure. Thus, polarisation resolved SHG (p-SHG) is used to acquire information about the orientation and degree of organisation of harmonophores, for example, to calculate the helical pitch angle of collagen fibrils [15] and the tilt angle of the helices relative to the fibril axis [16]. It has also been used to understand the differences between collagen types in tissues [17], and investigate the effect of mechanical stretching [18] on collagen disorder within tendon tissue. Collagen and its deposition/accumulation plays a critical role in many diseases especially fibrotic diseases such as liver fibrosis and idiopathic lung fibrosis [19] [20] [21]. The ability to resolve SHG-active structures at the nanoscale is vital as changes occur at the level of molecular-structures and assemblies such as the actin-myosin complex [22], microtubules [23] and fibrillar collagen [24–26] all of which are SHG-active. SHG has already proved to be valuable tool for biomedical and biophysical investigations however, sub-diffraction limited resolution has not been achieved preventing nanoscopic analysis.

Here we present the first demonstration of super-resolution SHG (SR-SHG) microscopy. We use PNJs generated using a microsphere to achieve SR-SHG and demonstrate a route to extended field of view of super-resolved SHG imaging. Most importantly, we find that polarisation properties of the excitation are maintained in a photonic nanojet. This unexpected finding crucially enabled polarisation resolved SR-SHG (pSR-SHG) that further enhances resolution. In an unprecedented and highly relevant application we show that nanoscopic anisotropy analysis enabled by pSR-SHG allows the detection of underlying nanoscale changes in collagen during the development of lung fibrosis. Overall, our work heralds the role that label-free super-resolution techniques can play to unravel underlying biological phenomena and improve our understanding of disease mechanisms.

## Results and Discussion

Achieving super-resolution SHG (SR-SHG) imaging can have applications in understanding nanoscale biology and evolution of SHG active materials such as in disease processes. Here, we used the photonic nanojet phenomena to achieve SR-SHG (Figure 1**A**). The SHG signals observed by exciting via a PNJ showed the expected 2^nd^ order dependence on power on imaging barium titanate nano-crystals (Supplementary Figure S 1**A-B**). In the context of SHG from single collagen fibres it has been shown that with higher excitation power, sensitivity to smaller fibre diameters is achieved [27]. We clearly also see that at higher power the signal to noise ratio improves in PNJ assisted SHG (Figure S 1**C-D**). Since SNR is inherently linked to resolution, further experiments were conducted at the highest excitation power possible (typically ~30 mW) within the damage threshold of the sample to obtain the highest resolution within the imaging conditions.

We acquired images of the same location of the sample without and with PNJ excitation of SHG to investigate the extent of improvement over conventional diffraction-limited SHG. A clear improvement in resolution is apparent in SHG signals from a lung tissue section (Figure 1**B,C**), that was verified to have an abundance of fibrillar collagen (Figure S3). We demonstrated that PNJ assisted SHG imaging allowed the distinction of individual fibres and their directionality, an observation that was not possible with conventional diffraction-limited imaging. A typical collagen fibre is between 12 and 500 nm (age dependent) in diameter as determined previously by electron microscopy [28]. The diffraction limited resolution of our microscope at the fundamental wavelength (800 nm) is ~330 nm. Hence, the mere fact that we qualitatively resolve collagen structures below this limit (Figure 1 **C**) indicates the super-resolution capability of PNJ excited SHG.

In order to quantitatively determine the resolution limit for SR-SHG a patterned silicon sample was fabricated of 5 individual cuts etched on a thin silicon membrane using a focussed ion beam. The width of each cut is 100 nm whilst the spacing between them is 125 nm (λ/6.4) verified by TEM (Figure 1**D inset**). Figure 1**D** shows an image acquired through a 14 μm diameter sphere to generate a PNJ. The relationship between size and magnification is shown in Figure S4; smaller spheres can achieve better resolution due to their narrower PNJ [6]. As shown in Figure 1**E** the slits (Figure 1**D**) can be well resolved satisfying the Rayleigh criterion (74% of peak). This shows that SR-SHG can be achieved with at least 125 nm resolution which is 1.68 times better than anything achieved in SHG microscopy [29]. In our experiments, the limitation was due to lack of available SHG-active standards and the fabricated sample exploited surface-SHG that is significantly weaker than bulk-SHG [30]. This could not be surmounted by applying higher laser power given the damage threshold for such thin samples. Hence, the actual achievable resolution with our PNJ-assisted SR-SHG method may be even higher.

Whilst super-resolution is clearly demonstrated with PNJ-assisted SHG, a potential limitation is the restriction of the field-of-view (FOV) given by one sphere. (Figure 1**C** shows 4 distinguishable stripes due to the location of the sphere on the sample as the field of view is limited by the sphere size.) We applied a simple multiplexing technique to allow rapid acquisition of multiple SR-SHG images using self-assembled ordered arrays of spheres (Figure 2 inset) to overcome this limitation. The magnification achieved is dependent on the size of the sphere used and the focal plane z-position (Figure S4). Therefore, if taken with an array of spheres, the image from neighbouring spheres can overlap and hence needs to be optimised. With a 60 µm diameter sphere a focal position of z=70 µm was found to be the optimum to maximise resolution whilst avoiding overlap of images (Figure 2). Although not demonstrated for SR-SHG, multiple spheres and image stitching has been used to reconstruct composite FOVs [31–33]. Our approach retains the advantage of simplicity whilst offering increased speed of data acquisition relative to single sphere approaches and most importantly is demonstrated for label-free laser-scanning nanoscopy (specifically SR-SHG).

**Figure 2:**
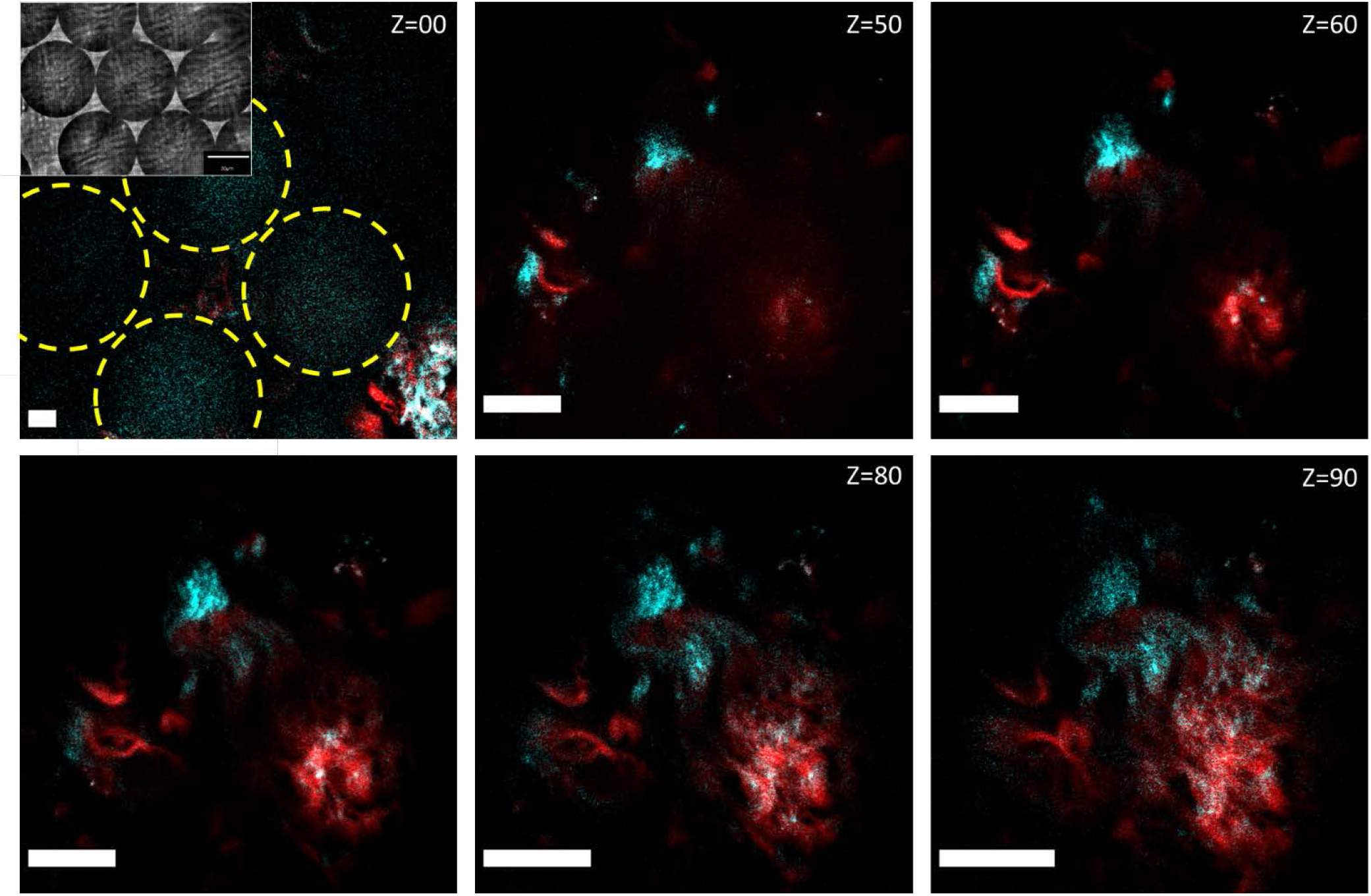
Arrayed spheres allow simultaneous acquisition of multiple images. Images shown of a group of spheres deposited on a lung tissue sample, cyan signal is SHG, red signal is two photon excited fluorescence. In the Z=00 image the locations of 4 neighbouring spheres can be seen indicated by yellow circles, 3 of these contribute to clear FOVs when imaging through the spheres. As the focal position is changed (Z is increased) the magnification increases as does the resolution. At Z = 70 µm magnification is maximised whilst ensuring non-overlapping FOVs, at greater values of Z overlap confounds image information from each sphere. Inset: Representative brightfield image showing 6 spheres packed together. Scale Bars 10 µm.

Having shown that SR-SHG is possible using the PNJ-approach and the FOV can be expanded, we sought to investigate ways to achieve ultra-high resolution and develop a technique that can be used for distinction at the nanoscale level. Hence we examined the property that in SHG, signals are dependent on the relative orientation of harmonophores vs the polarisation of the excitation [34] offering a measure of the disorder of a sample. We used a mouse tail tendon that contains a high density of aligned collagen fibres to demonstrate this sensitivity to polarisation (Figure 3). The polar plots (Figure 3**C**,**F**) showed the dumbbell shape as expected for polarisation dependent SHG signals [35]. However, this strong polarisation dependent behaviour of SHG observed even through a microsphere was intriguing. The experimental result suggests that polarisation is maintained despite complex propagation conditions (whispering gallery modes, near-fields etc) that are taking place in the microspheres and that are responsible for the tight focussing [2–9]. To verify this result, numerical simulations were performed to investigate the polarisation state of the photonic nanojet. Figure 3**G** and **H** shows the x and y components of the electric field (||E/E_0_||^2^) respectively when the sphere is illuminated with linearly polarised light in the x plane. The intensity of the y component in the PNJ is approximately 2 orders of magnitude smaller than the x component indicating that the polarisation state of the illumination is not significantly affected when focussed into a PNJ. These results indicate that it should be possible to perform super-resolution polarisation resolved SHG measurements.

**Figure 3:**
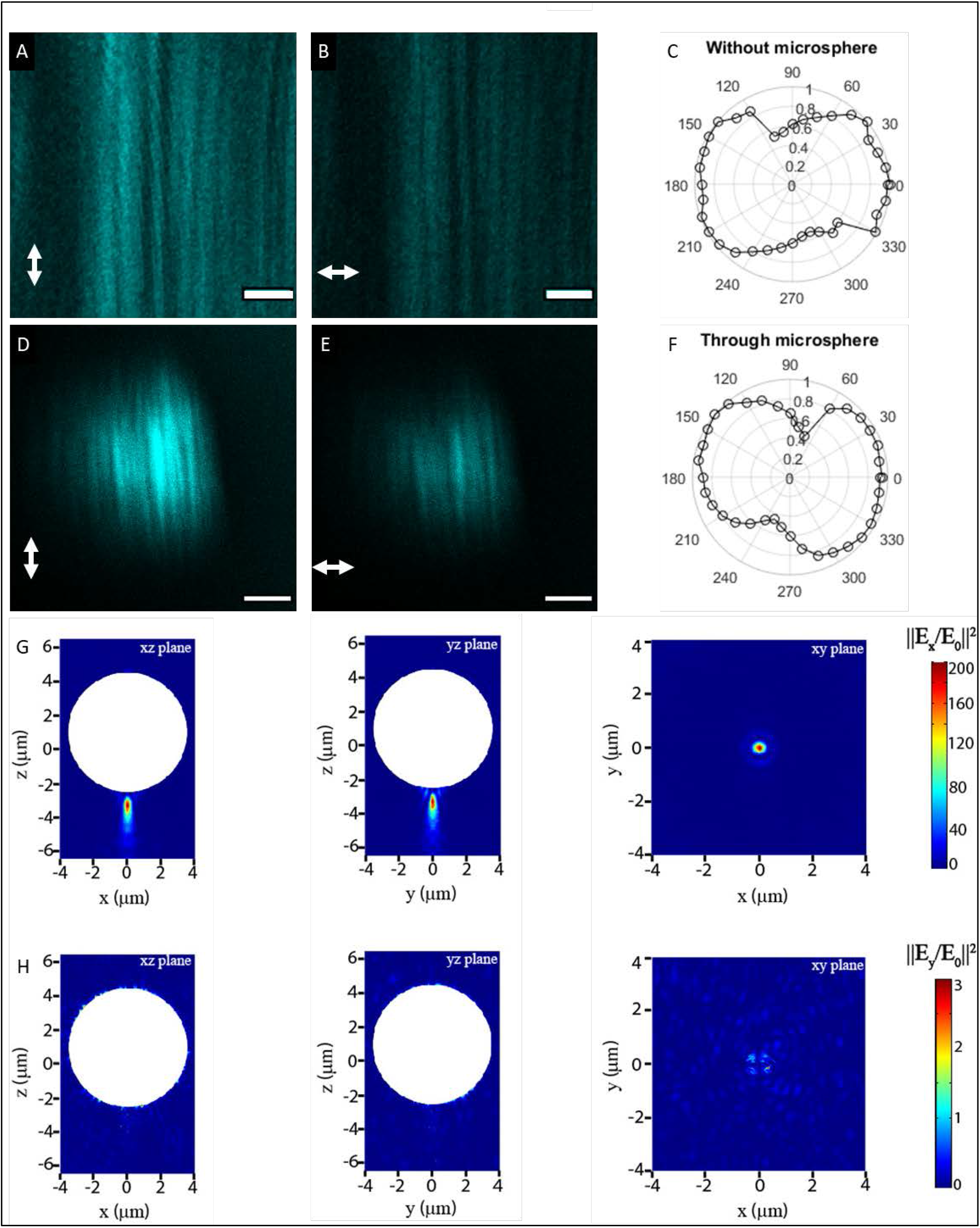
PNJs maintain the polarisation of light. **A** and **B**, images of collagen fibres in mouse tail tendon with the excitation polarisation aligned parallel and perpendicular to the fibre orientation respectively. **D** and **E**, the same situation as **A** and **B** with imaging performed using PNJs, in both cases the intensity is at a maximum when fibres and excitation polarisation are aligned and at a minimum when they are perpendicular. **C** and **F** show polar plots of signal intensity against excitation polarisation for the diffraction limited and PNJ cases respectively, the shape of the plots are the same showing that polarisation is maintained through microspheres. Scale bars 3 μm. **G**: Simulation results - Colour map plot of the x-component of the electric field intensity (||E/E0||2) in various planes. The incident light was x-polarised. The white shaded circular region represents the microsphere. The xz and yz corresponds to planes passing through the centre of the microsphere while the xy plane is plotted passing through the focal point of the generated PNJ, H: same as G but for the y component. The intensity of the y component is approximately 2 orders of magnitude smaller than the x component indicating that the polarisation state of the illumination is not significantly affected when focussed into a PNJ.

Polarisation dependent changes can be measured by polarisation anisotropy, *r* = (*I*_||_ − *I*_⊥_)/ (*I*_∥_ + 2*I*_⊥_), where *I*_||_ and *I*_⊥_ are the intensities recorded parallel or perpendicular to incident polarisation. Polarisation anisotropy can vary between −0.5 and 1 where 0 is completely random emission and 1 is emission completely aligned with the excitation polarisation. Pixel-wise calculation of anisotropy was performed to generate images (see Materials and methods) that show the spatial distribution of disorder within the collagen ultrastructure. We demonstrate that using the polarisation dependence of SHG through measurement of its anisotropy the spatial resolution can be increased. Figure 4**A** and **B** show the diffraction-limited while Figure 4**C** and **D** show the PNJ-assisted SHG intensity and corresponding polarisation anisotropy images. The same location taken from a sample containing collagen fibrils (lung cell culture model; see Materials and methods) was imaged. The polarisation anisotropy images reveal more structures than both the diffraction-limited and PNJ-assisted intensity images. Line profile plots show increased number of peaks and reveal hidden features in both diffraction-limited and PNJ-assisted SHG imaging. The anisotropy of SHG thus revealed the otherwise invisible fibres because polarisation is maintained by the PNJ. This demonstrates that polarisation resolved SHG, herein using the anisotropy offers improved super-resolution; we term these set of techniques as pSR-SHG. Polarisation has been used previously to improve the resolution of diffraction-limited fluorescence images [36,37] by improving the localisation accuracy of the dye molecules used for labelling biological samples. Here, however, we crucially show that polarisation anisotropy not only increases resolution of SHG but allows resolution of neighbouring nanoscale collagen fibrils improving both the sensitivity and the resolution even in PNJ-assisted (hence, super-resolved) SHG measurements whilst maintaining the benefits of label free imaging. The anisotropy of polarisation sensitive SHG diffraction-limited imaging itself is powerful and can be used to delineate different stages of ovarian cancer [38] and differentiate between different collagen isoforms [39]. However, in many cases the changes are much more subtle and occur at the nanoscale such as in idiopathic lung fibrosis [40]. Here we applied our simple super-resolved SHG imaging approach, particularly, pSR-SHG to see whether such nanoscale changes in collagen could be observed directly.

**Figure 4:**
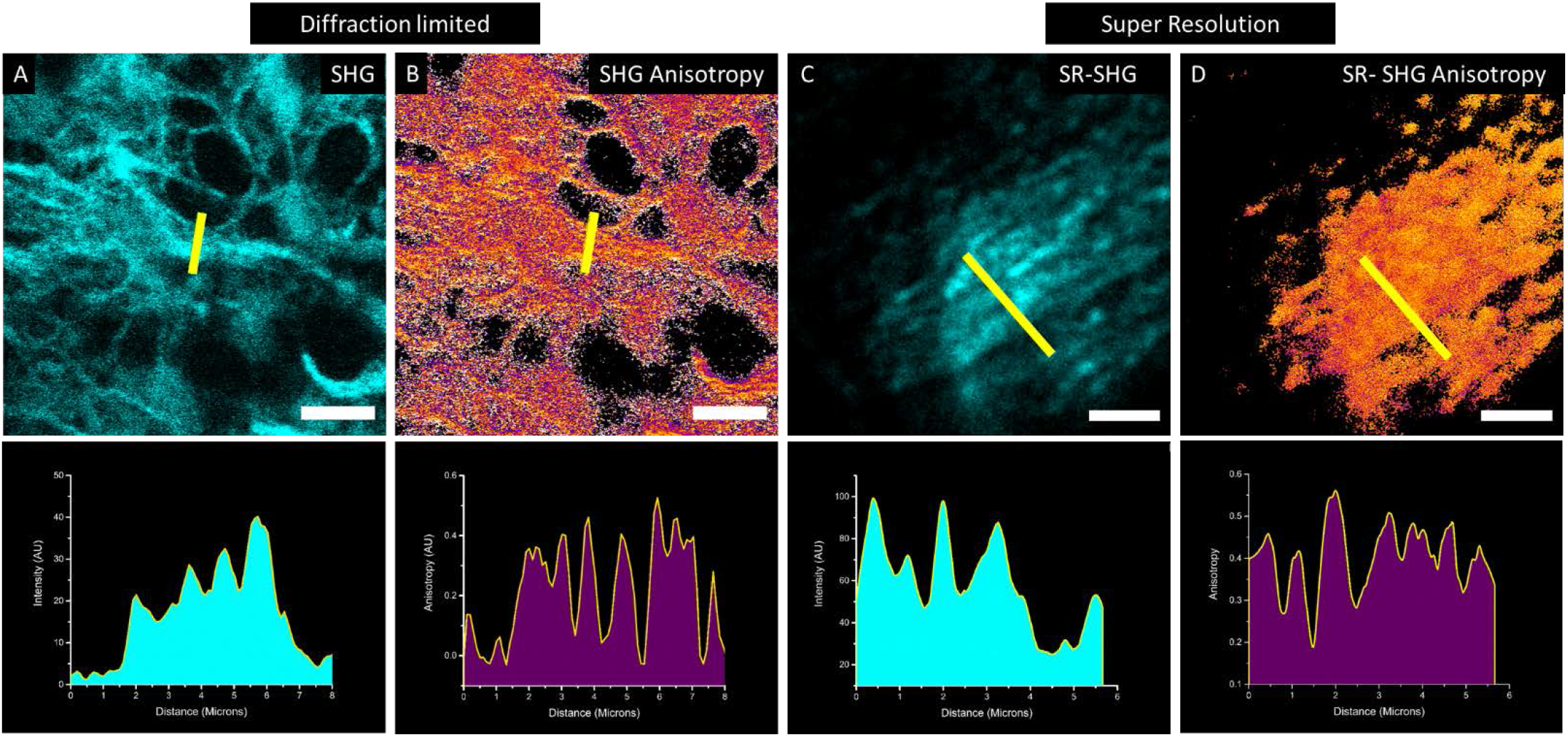
Polarisation anisotropy reveals hidden fibres that cannot be identified by intensity alone. **A** shows a diffraction limited SHG image of collagen fibrils and **B** shows the corresponding polarisation anisotropy image (scale bars 10 µm). Intensity plots taken through the yellow lines in **A** and **B** show that the polarisation anisotropy reveals more resolved peaks than the SHG intensity for the diffraction limited case. **C** and **D** show SR-SHG and corresponding polarisation anisotropy (pSR-SHG) images (scale bars 3 µm). Intensity plots taken through the yellow lines in **C** and **D** again show that the polarisation anisotropy reveals more resolved peaks than the intensity alone in SR-SHG as well.

Diffraction limited as well as PNJ-assisted SHG images of sections from a tissue spheroid model of the fibrotic lung disease, Idiopathic Pulmonary Fibrosis (IPF) were acquired (Figure 5). We have previously shown using transmission electron microscopy (Figure S5) that in this model the collagen ultrastructure changes significantly following treatment with lysyl-oxidase/ lysyl-oxidase-like (LOX/LOXL) inhibitor BAPN (ƅ-aminoproprionitrile) which inhibits cross-linking of collagen fibres [40]. Figure 5**A** shows diffraction limited SHG images for both the control and BAPN treated sections. We have previously demonstrated that the macroscopic collagen architecture becomes more disordered upon treatment with BAPN [40] and here this can be seen by the loss of the high intensity collagen ring at the edge of the sphere (Figure 5**A** red arrow). However, the central regions of the spheroids show very similar collagen architecture when viewed at the diffraction limit, which was further verified via image analysis of fibres using a number of parameters such as fibre length and width (Figure S6). The polarisation anisotropy measured from diffraction-limited images is unchanged (Figure 5**C**). SR-SHG images taken of the same spheroid samples also show that the collagen architecture is similar between the control and on BAPN treatment (Figure 5**B**, Figure S6). In contrast, pSR-SHG shows that there is a significant difference between the control and BAPN treated sample (Figure 5**D**). This is consistent with our previous observation using transmission electron microscopy of dysregulation of collagen fibril nano-structure throughout the spheroid model [40].

**Figure 5:**
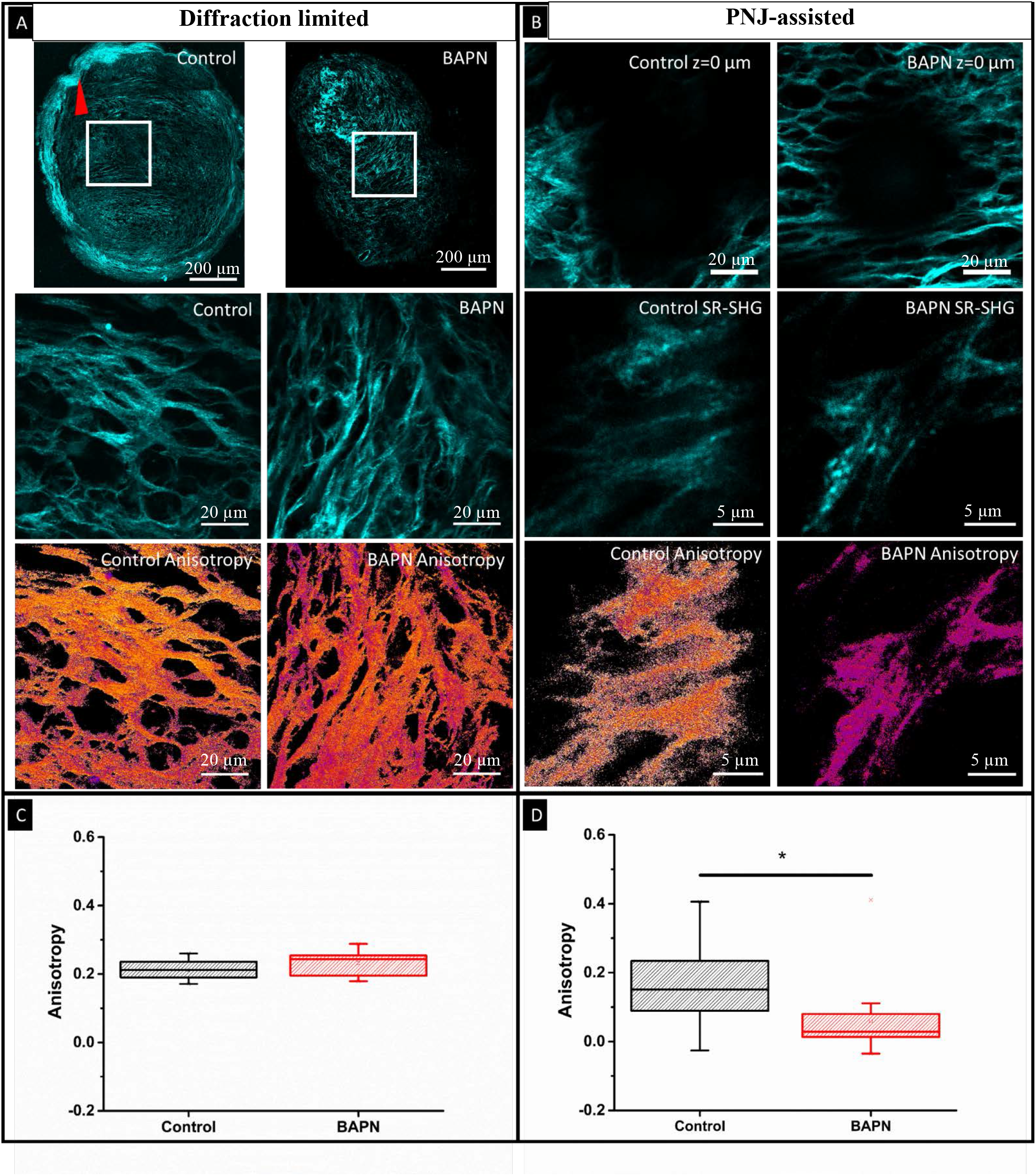
SR SHG polarisation anisotropy measurements reveal nanoscale differences in collagen ultra-structure under treatment with LOX/LOXL inhibitors. **A)** Diffraction limited imaging of Lung spheroids shows differing collagen morphology at the edge of the spheroid (loss of ordered collagen indicated by red arrowhead). Similar collagen morphology occurs within the central region of the spheroid between Control and treatment with LOX/LOXL inhibitor BAPN. The anisotropy images show that the measured anisotropy is unchanged between the treatments, which is quantified in **C** mean anisotropies being 0.2132 and 0.2322 for control and BAPN treated respectively(15 FOVs for control and BAPN),. **(B)** SR-SHG images show that the morphological similarities exist at a SR scale however; the SR-anisotropy **(D)** measurements highlight a significant difference in the mean anisotropy. Control: 0.1471(29 spheres) BAPN: 0.0600 (20 spheres). BAPN reduces the anisotropy. Significance tested using a one way ANOVA, P = 0.0131. **A**, scale bars = 200 µm in top row, 20 µm in middle and bottom rows, **B**, scale bars = 20 µm in top row, 5 µm in middle and bottom rows.

Currently imaging of such nanoscale changes in collagen is carried out by electron microscopy (EM) [41] and occasionally with atomic force microscopy (AFM) [42].While higher resolution is possible with these methods, none allow for the simplicity and hence can match the throughput of optical techniques. Using (p)SR-SHG methods demonstrated in this work, significantly high resolution can be achieved without the complexity of sample preparation or sophisticated equipment. Furthermore, in electron microscopy heavy metal stains need to be used to generate contrast [43] while for AFM analysis the collagen needs to be isolated [44] and analysis of native tissue is unrealistic.

## Conclusions

SR-SHG is able to improve the resolution of a conventional SHG microscope and facilitate label-free super-resolution imaging in a simple and cost effective way. In this work, we showed both qualitatively and quantitatively that PNJ-assisted SR-SHG can be achieved. We demonstrated a ~2.7X improvement over diffraction limited imaging. With the availability of suitable resolution standards for SHG imaging and using smaller spheres to generate PNJs even higher resolutions could be achieved. The key parameters affecting SR-SHG imaging were characterised and optimised including the effect of sphere size, and distance from the focal plan. We presented a way to increase FOV by using an array of self-assembled spheres. In a significant new finding we show that despite the tight focussing properties of microspheres the polarisation state is maintained in PNJs allowing pSR-SHG imaging, wherein polarisation anisotropy of SHG signals can be measured. pSR-SHG improved the resolution and sensitivity further over both diffraction limited as well over normal PNJ-assisted SR-SHG. In another significant result for a highly relevant application, we were able to observe nanoscale changes in collagen disorder in exemplar biological samples (lung tissue spheroid sections) that were invisible to diffraction-limited SHG imaging. Our (p)SR-SHG technique overcomes the issues of sample preparation and sophisticated instrumentation associated with AFM and EM techniques and is label-free. It can be performed on cells, native tissues (and their sections) directly making it highly suitable for biomedical research and clinical translation. (p)SR-SHG can provide unprecedented nanoscale insight into biological harmonophoric-structures such as collagen and thus has the potential to transform our understanding of a wide range of diseases and effects of treatments including in cancers and fibroses.

## Materials and methods

### SHG imaging system

Second harmonic generation imaging was performed on a home-built multiphoton imaging system consisting of a femtosecond pulsed laser (MaiTai, Spectra physics) that generates 120 fs pulses with a 80 MHz repetition rate, coupled into a Leica DMRB upright microscope stand. Laser scanning is performed using a pair of galvanometric mirrors (Cambridge Instruments).The beam is expanded using a telescope to overfill the objective back aperture (Leica 63x, 1.2 NA, water immersion) and is directed into the objective using a dichroic mirror (Semrock FF685-DI02-25X36). Signal is collected in the backwards direction and separated from the excitation laser by the same dichroic mirror, then cleaned using a short pass filter with a 694 nm cut-off. SHG was excited at 800 nm for all non-polarisation based images and at 808 nm for all polarisation based images. SHG separated from other signals using a dichroic mirror (Semrock, FF458-Di02) and detected after through a band pass filter centred at 400±20nm (Thorlabs, FB400-40). Detection was performed by focussing onto a PMT (Hamamatsu, H10722-01). For polarisation anisotropy measurements, an analyser (Thorlabs, LPVISE100-A) was placed directly in front of the PMT. The excitation polarisation was controlled using a combination of a half wave-plate (Thorlabs, WPH10M-808) to rotate the linearly polarised output of the laser and a quarter waveplate (Thorlabs, zero order 808nm) to pre-compensate for ellipticity induced by the scanning mirrors and excitation dichroic. The polarisation state of the fundamental beam at the sample was determined using a linear polariser (analyser) and power meter. For a given orientation of the analyser the extinction ratio was maximised at the linear polarisation of the fundamental deemed to be orthogonal to the analyser transmission axis.

**Scheme 1.**
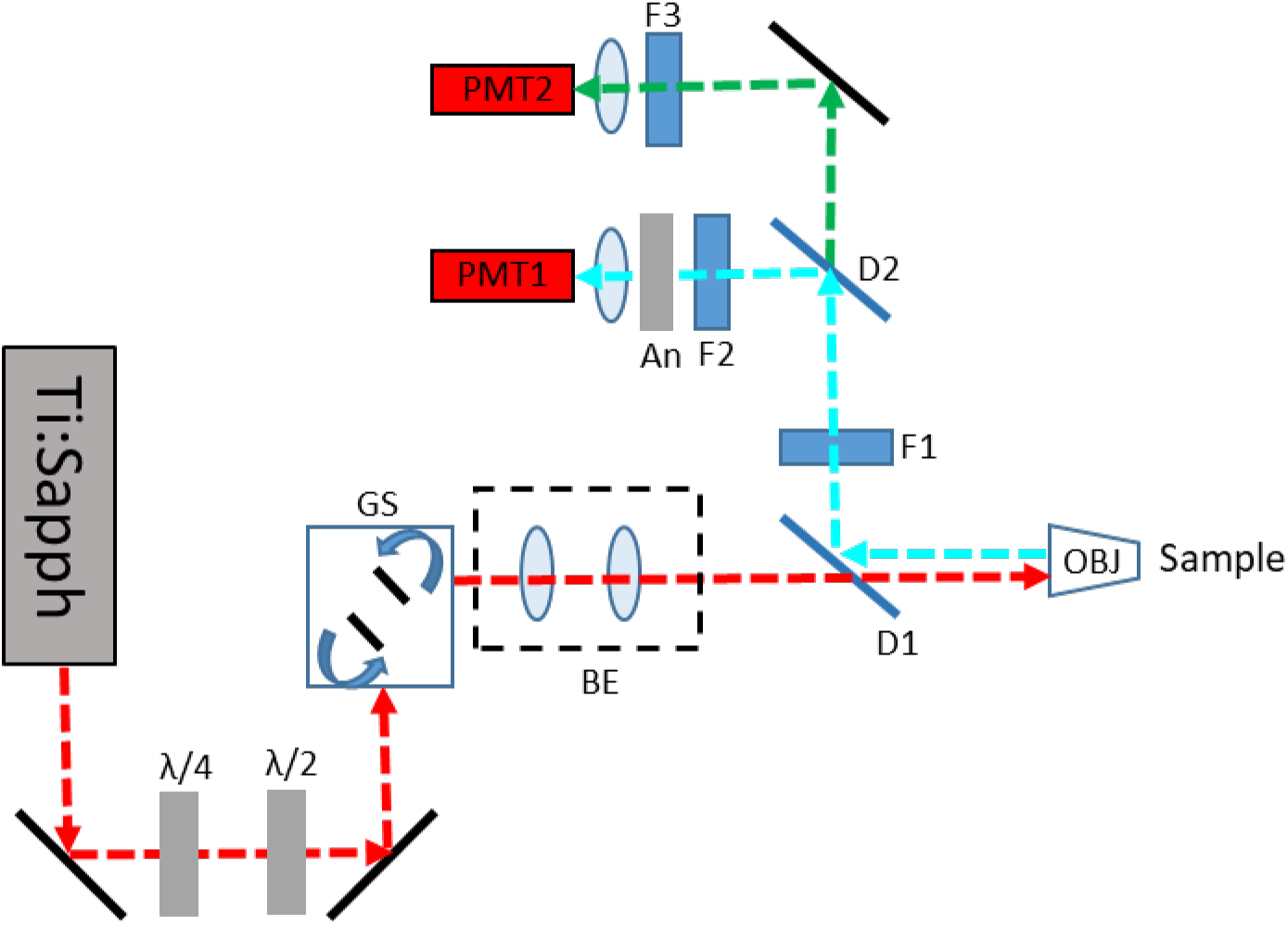
Schematic of the SHG imaging system. The linearly polarised output of a femtosecond pulsed laser (Ti:Sapph) is passed through a quarter(λ/4) and half waveplate(λ/2) to provide pre-compensation of induced ellipticity and to rotate the linear polarisation respectively. The beam is scanned by a pair of galvanometric mirrors, excitation is separated from emission via a dichroic mirror (D1) and signal cleaned using a short pass filter (D2). SHG signal is separated using a dichroic mirror (D2) and filtered (F2) before being focussed onto PMT1. For polarisation sensitive detection and analyser (An) was placed immediately after F2. Longer wavelength signals were detected on PMT 2 after filtering with an appropriate band pass filter (F3). GS: Galvo scanner, BE: Beam expander

### PNJ generation for imaging

Spheres made of Barium Titanate glass (BTG, refractive index of 1.9) were used. Their high refractive index contrast allowed them to be used with water (n=1.33) to form a photonic nanojet in water immersion medium with corresponding objectives commonly used for biological and multiphoton applications. Amorphous Barium titanate (BaTiO_3_) microspheres were purchased from Cospheric (Santa Barbara, CA, USA) in two size ranges of 5-22 μm and 53-63 μm. Spheres were suspended in water and a drop of the suspension used as the immersion medium for imaging.

### Barium Titanate Sample preparation

Barium Titanate nanocrystals <100nm specified size(Sigma-Aldrich)were suspended in ethanol, ultrasonicated for 10 minutes to break up aggregates, then a 10μL drop was placed onto a clean microscope slide and left to air dry.

### Mouse tendon sample preparation

All mouse experiments were carried out in accordance with the Animals (Scientific Procedures) Act 1986 set out by the UK Home Office. Female C57BL/6 mice aged between 4-6 months were culled via CO2 and cervical dislocation. The tail was removed using scissors and the tendon removed using a pair of forceps and was fixed in 4% paraformaldehyde for 6-8 hours. They were then washed in Phosphate Buffered Saline (PBS) immediately before being mounted on a microscope slide and sealed under a coverslip (#1.5; 0.17 mm thick) before imaging.

### Lung tissue

Dewaxed formalin-fixed paraffin-embedded human lung tissue sections (5 μm) were from tissue surplus to clinical requirements from lung biopsies of patients with a subsequent diagnosis of idiopathic pulmonary fibrosis (IPF) under the approval of the Southampton and South West Hampshire and the Mid and South Buckinghamshire Local Research Ethics Committees (ref 07/H0607/73). All subjects gave written informed consent.

### Spheroid sample preparation

Details of spheroid growth have been described in detail previously [40]. Briefly: Primary lung fibroblasts from a patient with IPF were seeded in Transwell inserts under normal conditions for 24 hours before the media was replaced with that containing BAPN or control conditions. After 6 weeks of growth under these conditions the spheroids were harvested, fixed in 4% paraformaldehyde, paraffin embedded, and 5µm sections cut ready for imaging. For transmission electron microscopy(TEM) spheroids were fixed in 3% glutaraldehyde in 0.1M caco-dylate buffer at pH 7.4.

### Transmission electron microscopy

As previously described [40] spheroids were post-fixed sequentially in osmium/ferocyanide fixative thiocarbohydrazide solution, osmium tetroxide, uranyle acetate and Walton’s lead aspartate solution before dehydration in graded ethanol and acetonitrile. Samples were embedded in Spurr resin and 100nm ultra-thin sections visualised using an FEI Tecnai 12 transmission electron microscope (FEI Company, Hillsboro,OR,USA).

### SHG Resolution standard fabrication

Silicon membranes were commercially sourced (Norcada, Inc.), 200 nm thick polycrystalline silicon membrane 1×1mm, 200µm Silicon frame 5×5mm. FIB was performed on an FEI Helios NanoLab 600 FIB system. Each cut was formed by using an ion beam of Ga+ ions, with an acceleration voltage of 30 keV and beam current of 28 pA.

### Simulations

Numerical simulations were carried out using COMSOL Multiphysics 5.4, a FEM (finite element method) based numerical software to investigate light propagation through the barium titanate microsphere (refractive index was assumed to be nms =1.9). We adopted full field formalism with periodic boundary conditions (Floquet) applied on all the four sides of the computational geometry assuming an infinite 2D periodic structure in the x-y plane, solving the wave equation in a three-dimensional model. The incident beam was assumed to be a plane wave, linearly polarised along x axis propagating from the front side of the microsphere ie along the z direction and the excitation wavelength was set at 800 nm. The size of the microsphere was taken to be 7 µm and was positioned at the centre of the computational window and also assumed to be embedded in water medium (n = 1.33). The relative permeability was taken to be µr =1. The entire structure was divided into domains and sub-domains, and each domain was meshed using free tetrahedral meshing of maximum element size 6 elements per wavelength outside the microsphere and extremely fine meshing within the sphere. Direct solver method was adopted to solve for the wave equation.

### Calculation of Polarisation anisotropy

Polarisation anisotropy images were created using a home written Fiji macro in which the key steps are as follows: A difference image was generated by (I_par_ – I_perp_), this was divided by a total intensity image (I_par_ + 2I_perp_). The total intensity image was used to select the region of the image containing signal and for display all pixels outside of this region were set to 0. Only the regions containing signal were used for further anisotropy analysis.

### Collagen fibre analysis

Collagen fibre analysis was performed using CT-FIRE V1.3. SHG intensity images were converted to 8bit format before being batch processed using default parameters. Four parameters were measured to describe fibre morphology, width, length, straightness and angle.

## List of abbreviations

SHG: Second Harmonic Generation
SR: Super-resolution
BTG: Barium Titanate Glass
PNJ: Photonic Nanojet
PSF: Point Spread Function

