## Supporting information for "Super-resolved polarisation-enhanced second harmonic generation for direct imaging of nanoscale changes in collagen architecture"

### 1. Confirmation of SHG signals excited via photonic nanojet and resolution improvement with increased power

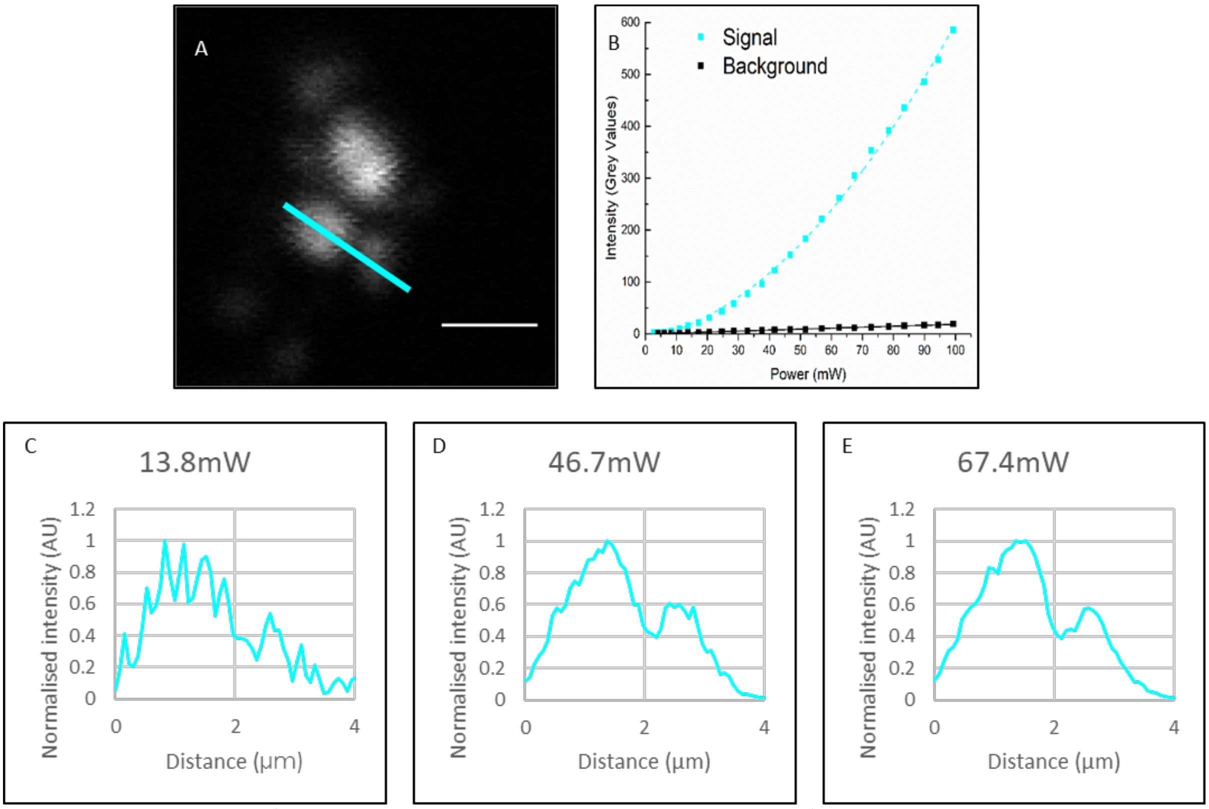


**Figure S1: Characterisation of the SHG response when using PNJs**. **A** an example image of Barium Titanate nanocrystals imaged through a microsphere at increasing incident powers. Scale bar 2µm. **B** Plots of signal and background against incident power, the signal shows a quadratic relationship to power whereas the background increases linearly confirming the signal as SHG. **C,D,E** Intensity plots taken across the cyan line marked in **A** as power is increased **(C→E)** the signal to noise ratio increases and the peaks become more easily resolved. Increasing the power improves resolution.

### 2. Improvement in resolution on SHG-active standard samples

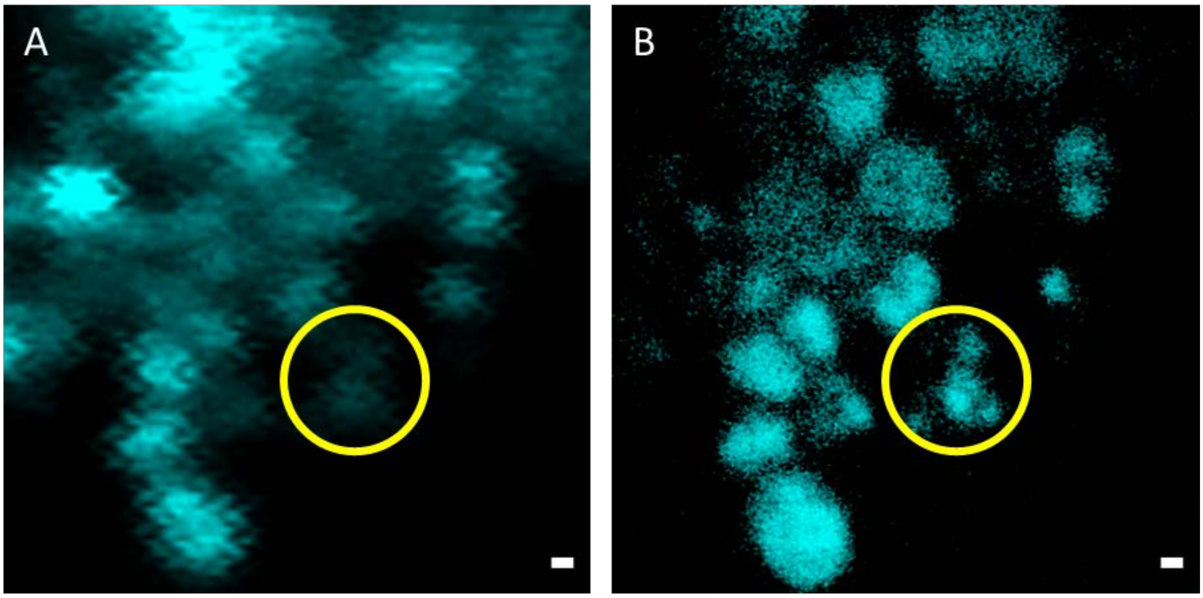


**Figure S2. PNJs improve the resolution of a Second Harmonic microscope**. **A:** A diffraction limited SHG image of $BaTiO_3$ nanocrystals and **B** an SR-SHG image of the same location. Scale Bar 500 nm.

### 3. Verification of presence/abundance of collagen in tissue samples investigated in this study

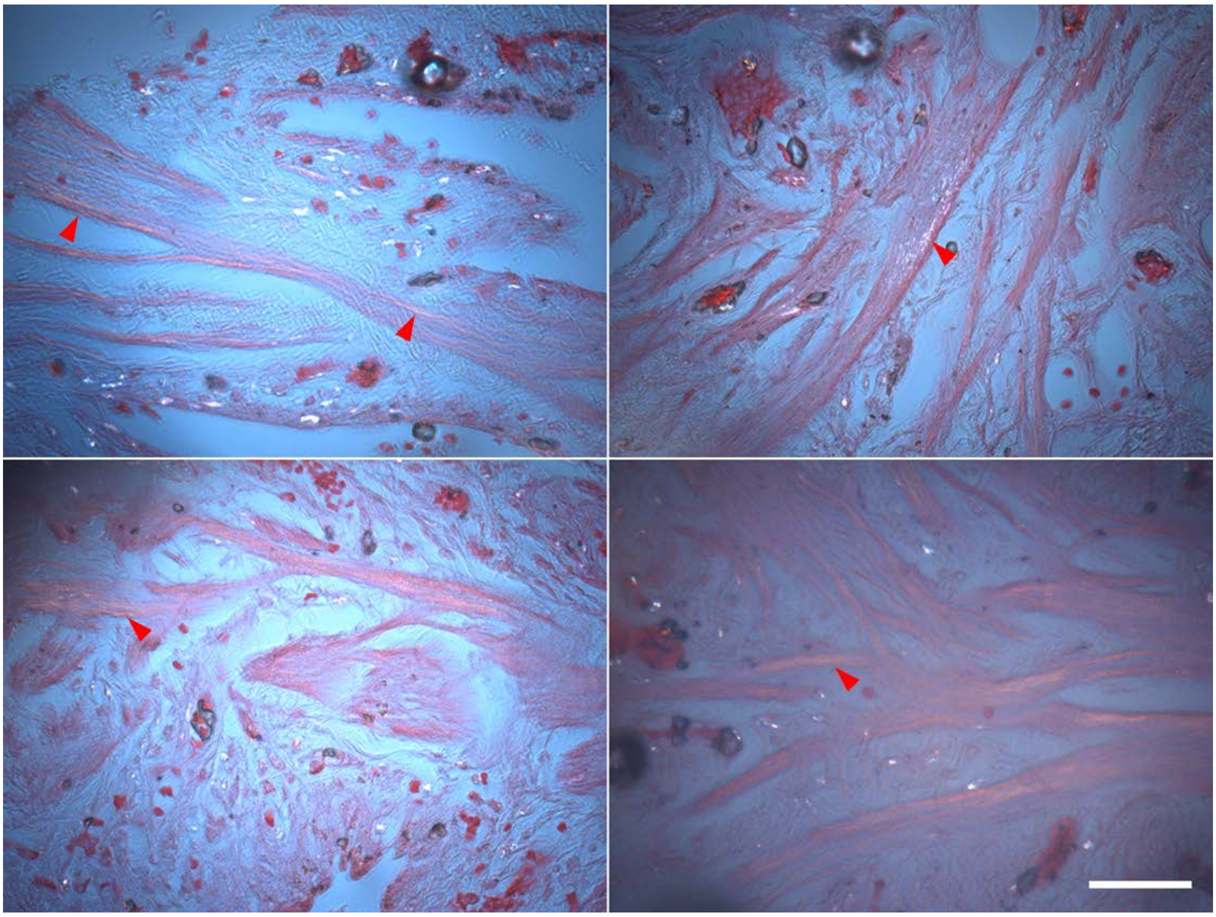

**Figure S3 Confirmation of collagen abundance in the tissue samples studied.** Example images taken of the same lung tissue sample (with a parallel section) showing the distribution of collagen stained with PicroSirius Red and imaged using polarised light microscopy. Collagen fibres appear as gold/white on a pink and blue background, red arrows indicate areas of high collagen concentration Scale bar 50µm.

### 4. Optimization of parameters for PNJ-assisted SR-SHG

Previous studies have shown that the degree of magnification and the improvement in resolution generated by a microsphere are dependent on the sphere size [45] and the relative z position [46] of the objective focus and the sphere-sample contact. To characterise this for SHG imaging a patterned sample of known dimensions was generated from a 200 nm thick silicon film. A series of slits were cut into the film using focussed ion beam (FIB) milling, the interface between the silicon and the immersion water that filled the slit provided the lack of inversion symmetry required to be SHG active. PNJ behaviour consistent with previously published results [9,47] was observed. Shorter and narrower PNJs are formed by smaller spheres [48] [6] however, they have a focal range closer to the sphere-sample contact. For thicker samples, out of focus excitation not contributing to the PNJ creates a strong background over which the signal is not observable. Therefore, the relationship between imaging range and sample thickness must be considered when selecting a sphere size to use. Sphere size also affects the achievable resolution in SR-SHG with smaller spheres providing better resolution due to the smaller photonic nanojet [6] but with a smaller field of view. Based on all these considerations, and given that tighter PNJs are expected of smaller spheres we chose 14 µm spheres at the z-position z+20µm for highest magnification for establishing the limit of resolution of SR-SHG while for tissue imaging we chose 60 µm spheres to increase FOV whilst maintaining sub-diffraction limited resolution.

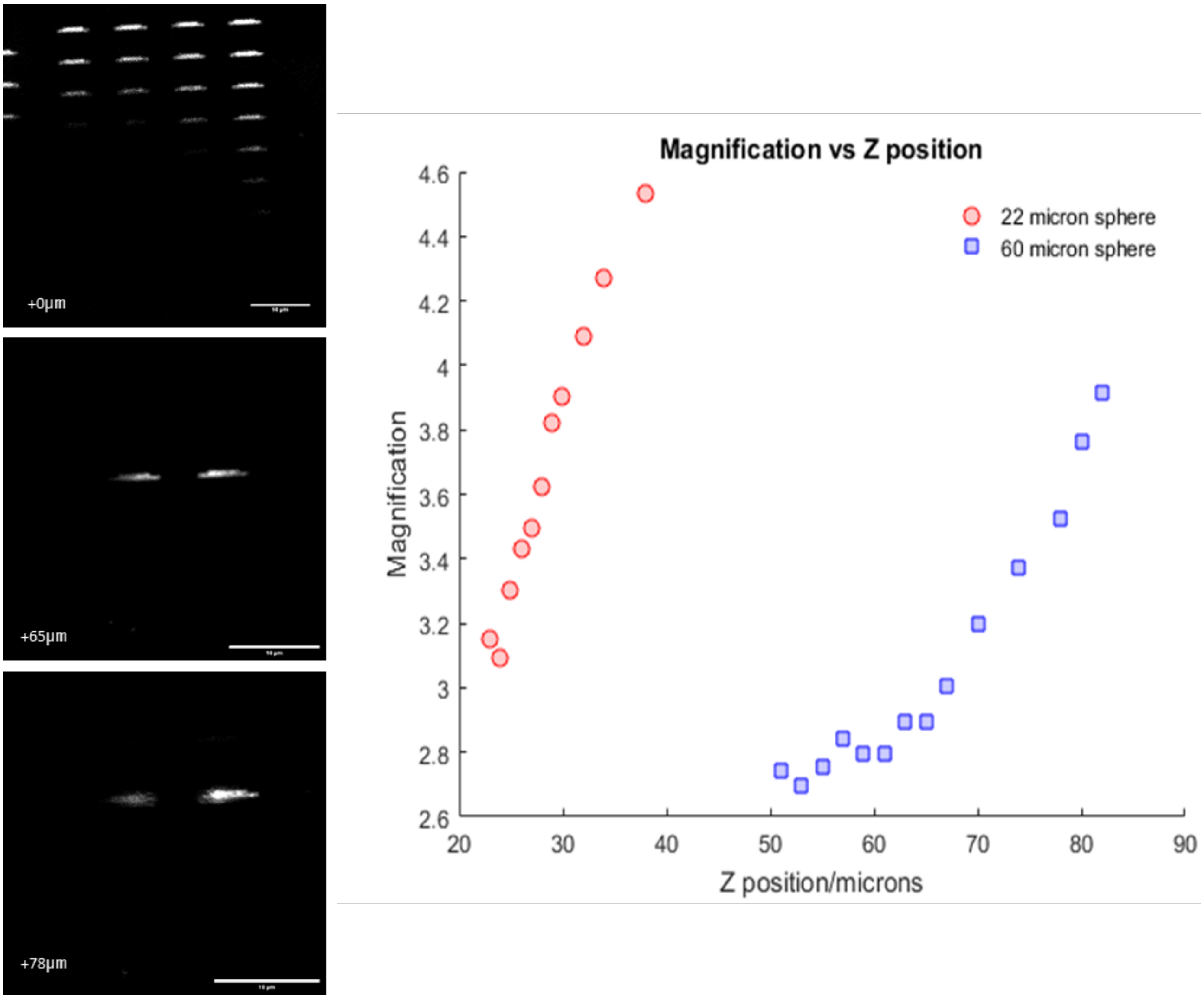


**Figure S4 Magnification dependence with PNJs.** The degree of magnification by microspheres depends on the relative position of the objective focal plane and the sphere sample contact (Z position) and the size of the sphere. 3 example images taken through a 60µm sphere. The top image is taken focussed at the sample surface, the sample contains a grid of 4 x 8 slit features, and the dark region indicates the location of the sphere. The images taken at z+65 µm and z+78 µm show that the image generated through the sphere increases in magnification as z increases. The z+78 µm image also shows that the image starts to become defocussed as z increases past the focal range, Images were taken through microspheres of 2 different sizes, 22 µm and 60 µm, The z-ranges for obtaining magnified images were 50-85 µm for the larger spheres and 20-40 µm for the smaller spheres. The graph shows that smaller size spheres produce a greater magnification than the larger spheres but over a narrower focal range. The focal range of smaller spheres occurs closer to the sphere than that of larger spheres. Scale bar 10 µm.

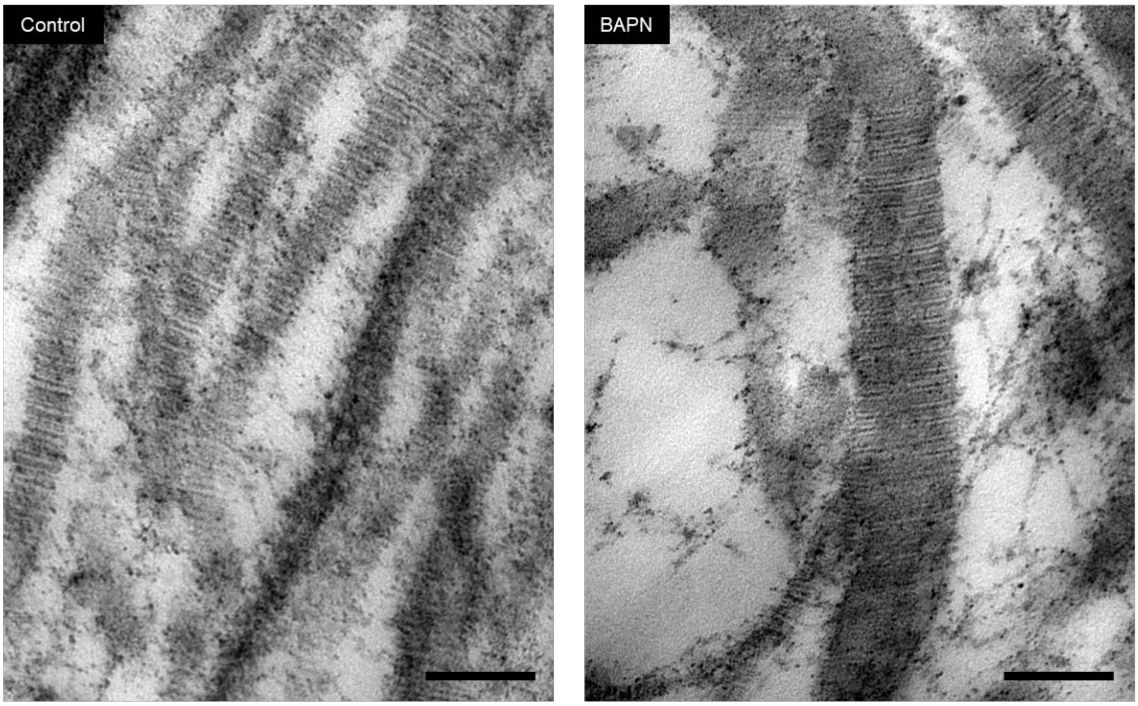


**Figure S5: TEM verification of changes in collagen architecture on treatment with BAPN.** IPF fibroblasts were grown for 6 weeks in the 3D in vitro model of fibrosis with vehicle control, or 1 mM BAPN, as indicated. Transmission electron microscopy images of collagen fibrils in longitudinal cross-section identifies that with BAPN there is an increase in fibril diameter and a marked dysregulation of fibril structure including irregular profiles. Scale bar 100 nm.

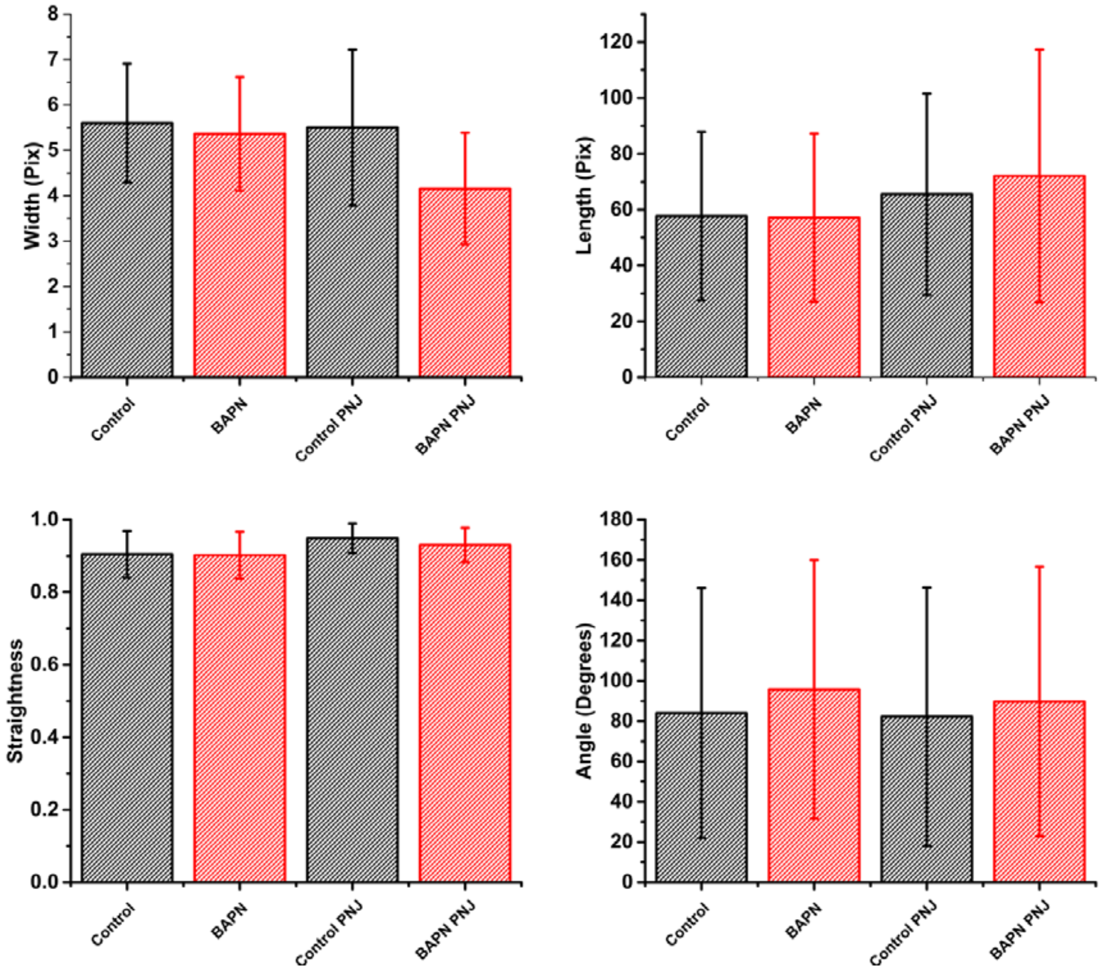


**Figure S 6: Conventional analysis of collagen fibres.** Image analysis of fibre parameters shows that there is no observable difference between the central regions of Control and BAPN treated spheroids. CT-FIRE was used for the analysis. Four parameters were measured to describe fibre morphology, width, length, straightness and angle, for each of these parameters there is no significant distance between control and BAPN treated case. A total of 15 FOVs were tested for Control and BAPN diffraction limited case and 20 FOVs for PNJ assisted images. Error bars = 1 standard deviation from the mean.
